# Effects of emotion and semantic relatedness on recognition memory: Behavioral and electrophysiological evidence

**DOI:** 10.1101/2020.02.28.969659

**Authors:** Meng Han, Bingcan Li, Chunyan Guo, Roni Tibon

**Author notes:** Corresponding authors: Roni Tibon School of Psychology, University of Nottingham University Park, The University of Nottingham, Nottingham, NG7 2RD, Chunyan Guo School of Psychology, Capital Normal University No.23 Baiduizijia, Fuwaidajie St. Haidian District, Beijing, 100048 PR China.

## Abstract

Some aspects of our memory are enhanced by emotion, whereas other can be unaffected or even hindered. Previous studies report impaired associative memory of emotional content, an effect termed associative “emotional interference”. The current study used EEG and an associative recognition paradigm to investigate the cognitive and neural mechanisms associated with this effect. In two experiments, participants studied negative and neutral stimulus-pairs that were either semantically related or unrelated. In Experiment 1 emotions were relevant to the encoding task (valence judgment) whereas in Experiment 2 emotions were irrelevant (familiarity judgment). In a subsequent associative recognition test, EEG was recorded while participants discriminated between intact, rearranged, and new pairs. An associative emotional interference effect was observed in both experiments, but was attenuated for semantically related pairs in Experiment 1, where valence was relevant to the task. Moreover, a modulation of an early associative memory ERP component (300-550 ms) occurred for negative pairs when valence was task-relevant (Experiment 1), but for semantically related pairs when valence was irrelevant (Experiment 2). A later ERP component (550-800 ms) showed a more general pattern, and was observed in all experimental conditions. These results suggest that both valence and semantic relations can act as an organizing principle that promotes associative binding. Their ability to contribute to successful retrieval depends on specific task demands.

## Introduction

Convergent evidence indicate that emotions influence memory, but the ways in which they operate and manifest require further specification (Easterbrook, 1959; Kensinger, 2009; Mackay et al., 2004; Mather & Sutherland, 2011). The term *emotionally enhanced memory* (EEM) refers to the common finding that emotional events are remembered better than neutral ones; a phenomenon that had been demonstrated for memory of emotional words, pictures, and stories (e.g., Bradley et al., 1992, with pictorial stimuli; Cahill & McGaugh, 1995, with stories; and LaBar & Phelps, 1998, with taboo words). According to one suggestion (Kensinger, 2009), this advantage for emotional content, and negative content in particular, arises from increased engagement of sensory processes, leading to focused attention on intrinsic details and to focal enhancement of the negative event. A similar account is the object-based framework (see also arousal-biased competition; Mather & Sutherland, 2011), which posits that emotional objects attract attention that enhances within-item binding of their constituent features (Mather, 2007). Accordingly, studies have shown that emotion facilitates memory for within-object memory binding (Kensinger, 2009; Mather & Nesmith, 2008; Mather & Sutherland, 2011; Nashiro & Mather, 2011; Schmidt et al., 2011; Steinmetz et al., 2015).

In contrast to the facilitative effects of emotion on within-item binding, inter-item associative memory (i.e., remembering that two or more items were encoded together) for emotional content is often hindered, an effect termed associative “emotional interference” (Madan et al., 2012; Mao et al., 2015; Mather & Knight, 2008; Pierce & Kensinger, 2011; Rimmele et al., 2011). One method of accessing associative episodic memory is recognition, i.e., the judgment that currently presented items were previously experienced together in a specific episodic context. In an associative recognition task, during an initial study phase, participants memorize pairs of stimuli. Then, at a subsequent test phase, they are asked to discriminate between intact (studied pairs), rearranged (studied items with new combination), and new pairs (of unstudied items). Thus, successful performance require memory for the specific pairings of the stimuli that were encoded. Pierce and Kensinger (2011) used this task to investigate the effect of emotion on associative memory. In their version of the task, studied pairs included either emotionally arousing or neutral words-pairs. Whereas for intact pairs accuracy did not differ, emotionally arousing rearranged word-pairs (and negative ones in particular) were less likely to be correctly identified as rearranged relative to neutral word-pairs. This interaction between emotional effects and retrieval conditions points to the multiplicity of processes involved in associative recognition, and their varying sensitivity to the influence of emotional content.

The prevalent dual-process model of recognition memory suggests that recognition involves two separable processes: familiarity and recollection. Familiarity is a general feeling of encountering something or someone which was previously seen, without memory of additional details from that initial encounter, whereas recollection involves the conscious retrieval of additional contextual details (for review see Yonelinas, 2002). Amongst other evidence, this distinction is supported by a large corpus of event-related potential (ERP) studies, revealing two distinct components which are differentially engaged in recognition judgments. The first component is considered to be the electrophysiological index of familiarity. It onsets ∼300-500 ms post-stimuli presentation, elicits greater negativity for new vs. old items, and is prominent over mid-frontal locations. The second component is considered to be an electrophysiological index of recollection. It onsets ∼500-800 ms post-stimuli presentation, elicits greater positivity to old vs. new stimuli, and is usually (though not always) prominent over parietal locations (reviewed by Curran, 2000; Diana et al., 2007; Rugg & Curran, 2007).

Previous studies investigating the effects of emotion on these ERP components yielded mixed results. Several studies used a context effect paradigm, in which neutral stimuli were encoded in the context of a neutral/negative sentences (Maratos & Rugg, 2001) or scenes (Smith et al., 2004; Ventura-Bort et al., 2016), and were subsequently discriminated from new stimuli via an old/new recognition judgment task. In one report (Ventura-Bort et al., 2016), the valence of the encoding context modulated the early and late old/new ERP effects. In another (Maratos & Rugg, 2001), a similar modulation was observed but only in one (out of two) of the experiments reported in the study. Conversely, in a third report (Smith et al., 2004), neither the early old/new component, nor the late one, were modulated by the valence of the context, but instead additional early ERP components were elicited for items encoded in emotional contexts. In addition, several studies that employed a source-memory paradigm, in which participants are explicitly asked about the encoding context, reported increased modulation of the old/new ERP effects for emotional contexts (Minor & Herzmann, 2019; Newsome et al., 2012), whereas others reported an opposite pattern (Mao et al., 2015). Finally, using subjective measures of familiarity and recollection (via remember/know or confidence judgments), several studies reported stronger recollection for emotional vs. neutral stimuli, accompanied by larger ERP old/new effects for the former (Schaefer et al., 2011; Ventura-Bort et al., 2016; Weymar et al., 2010; Weymar et al., 2011). In the current study, we utilized an associative recognition task, together with the well-established distinction between familiarity and recollection, and the putative electrophysiological markers corresponding to these processes, to aid our understanding of the mechanisms that drive emotional effects on associative memory.

Furthermore, the current study sought to examine how pre-existing semantic relations between episodically associated stimuli interact with emotional effects on associative memory. Semantic relatedness might affect memory two different ways. First, congruent stimuli combinations, such as related (vs. unrelated) pictures, support high Levels of Processing (LOP; Craik & Lockhart, 1972), yielding rich and elaborated encoding. In turn, during retrieval, this elaborated encoding supports high levels of recollection (summarized by Craik, 2002). Second, stimuli that share pre-existing relations can be subjected to unitization—an encoding process by which several discrete items are perceived and encoded as one single unit (reviewed by Mecklinger & Jäger, 2009; Yonelinas et al., 2010)—and to be subsequently retrieved via familiarity (e.g., Li et al., 2019; Tibon et al., 2014; Zheng et al., 2015). This latter suggestion is rooted in the idea that instead of creating inter-item links between the items, unitization results in a joint representation of the encoded items (that is, a single item, rather than several linked items). Importantly, unitization is not only enabled via explicit instructions to process pairs of memoranda as a single unit, but can also be driven by stimulus- or associative-properties of the encoded items (e.g., Tibon, Gronau, et al., 2014). This notion had been demonstrated in several ERP studies, in which the putative electrophysiological correlate of familiarity—the early old/new effect—was observed for episodically formed associations between semantically related stimuli, but not for episodic associations between semantically unrelated stimuli (Li, et al., 2019; Rhodes & Donaldson, 2007; Tibon et al., 2014; Tibon, Gronau, et al., 2014; Zheng et al., 2015).

The possibility that unitization might correspond to semantic relatedness effects is especially intriguing given the emotional enhancement effect of within-object memory binding (Mather & Sutherland, 2011). More specifically, it has been argued that when items are unitized, rather than relying on inter-item binding, their relations are supported by links that are qualitatively similar to within-object links (e.g., Mayes et al., 2007; Tibon et al., 2018). Therefore, unitized associations might be subjected to the enhancing effects of emotions on item memory. In other words, if emotions benefit within-object binding, then unitization of emotional stimuli might alleviate the negative effects of emotion on associative memory. Indeed, previous studies showed that when several items were integrated together (e.g., by creating a mental image of the words comprising a pair interacting with each other. For example, for the word pair “flies-booger”: imagining flies stuck in booger), associative emotional interference can disappear or even transform into mnemonic benefits (e.g., Guillet & Arndt, 2009; Murray & Kensinger, 2012).

Finally, the current study examined whether emotional and relatedness effects on associative memory depend on the relevancy of emotions to the encoding task. This point is further elaborated below, but in short, previous studies suggested that emotional content effects on processing and behavior often emerge when emotions are relevant to the task that is performed (e.g., Engen et al., 2017; Huang et al., 2008; Pessoa, 2009; Stein et al., 2009; Wei et al., 2015), though it is unclear whether this also applies to memory. Accordingly, in the present study we conducted two experiments, in which emotions were either relevant (Experiment 1) or irrelevant (Experiment 2) to the encoding task. In both experiments, an associative memory paradigm was used in order to examine the effects of valence (negative, neutral) and relatedness (related, unrelated) of memoranda on associative recognition.

## Experiment 1

In experiment 1, participants memorized and retrieved picture-picture pairs while EEG was recorded. Half the pairs were of negative objects (e.g., gun, spider) and half of neutral objects (e.g., bunny, traffic light). In addition, half the pairs contained objects that were semantically related (e.g., bunny-carrot), whereas half contained objects with no pre-existing semantic relations (e.g., spider-bullet). During study, participants performed a valence judgment task, and at test they were asked to discriminate between intact, rearranged and new pairs.

We predicted an overall reduction in associative memory accuracy for negative vs. neutral pairs due to associative emotional interference. Importantly, possible moderation of this effect by semantic relatedness will crucially depend on the mnemonic processes promoted by semantic relations: either unitization or LOP. More specifically, if semantic relations promote unitization, then associative retrieval of semantically related pairs would benefit from within-item binding. In this case, we would expect an attenuation of the associative emotional interference effect for negative pairs that are semantically related (compared to unrelated). We therefore predicted that associative memory for negative-related pairs would be similar (if not comparable) to memory for neutral-related pairs. We further predicted that this attenuation will be accompanied by a modulation of the early ERP component—the electrophysiological marker of familiarity—for related but not for unrelated pairs. Thus, the early associative memory ERP effect was expected to be greater for related pairs than for unrelated pairs. If, on the other hand, relatedness increases LOP, then we would not expect the associative emotional interference effect to be attenuated for related pairs (indeed, we might even observe the opposite pattern). Moreover, we would expect increased modulation of the late ERP—the marker of recollection—for related compared to unrelated pairs.

### Method

#### Participants

To determine the required sample size, we first extracted the behavioral effect size obtained by Pierce & Kensinger (2011; N = 32, Cohen’s f = 0.42), who employed a similar experimental design to the one used in the current study. Next, we extracted the associative memory ERP effects (for the early and late ERP components) from multiple studies that used an associative memory paradigm (Bader & Mecklinger, 2017; Kamp et al., 2016; Li et al., 2017; 2019; Rhodes & Donaldson, 2009; Tibon, et al., 2014; Zhao et al., 2020; Zheng et al., 2015). The effect sizes for the associative memory component reported in these studies were all medium-large, ranging from Cohen’s f = 0.3 to 10.7 (N ranging between 17 and 46). Nevertheless, to avoid an overinflated estimation of effect size, we set f = 0.25 as a lower, more conservative value. Based on this effect size, we estimated that power > .8 (α = .01) would require at least 39 participants (actual power = .81), and therefore recruited 47 participants for the study.

Forty-seven healthy, right-handed native Chinese speakers (30 females; mean age 22.4 ± 2 years) from Capital Normal University participated in the experiment and were paid ¥30 per hour. All participants had normal or corrected-to-normal vision, were prescreened for history of neurological or psychiatric disorders, learning disorders, head injury or psychotropic drug use. Informed consent, approved by the Capital Normal University Institutional Review Board, was collected from each participant. Data from five participants were discarded, including one participant with very poor task performance (associative Pr < 0), and four participants with insufficient number of artifact-free ERP trials in one or more experimental conditions (N < 16). Our final sample therefore included 42 participants (27 females; mean age 22.4 ± 2 years).

#### Stimuli

Our initial stimuli dataset included 1469 object pictures of animals, food, equipment, tools, appliances, etc., from the Hemera Photo-Objects Collection (Hemera Photo Objects, Gatineau, Quebec, Canada), the International Affective Pictures System (Lang et al., 2008) and free internet sources. Colored objects were matched for size, luminance and contrast by using adjustment curve in Adobe Photoshop 8.0, and were presented at the center of the picture on a gray background (RGB: 150; see Fig. 1 for examples).

**Figure 1.**
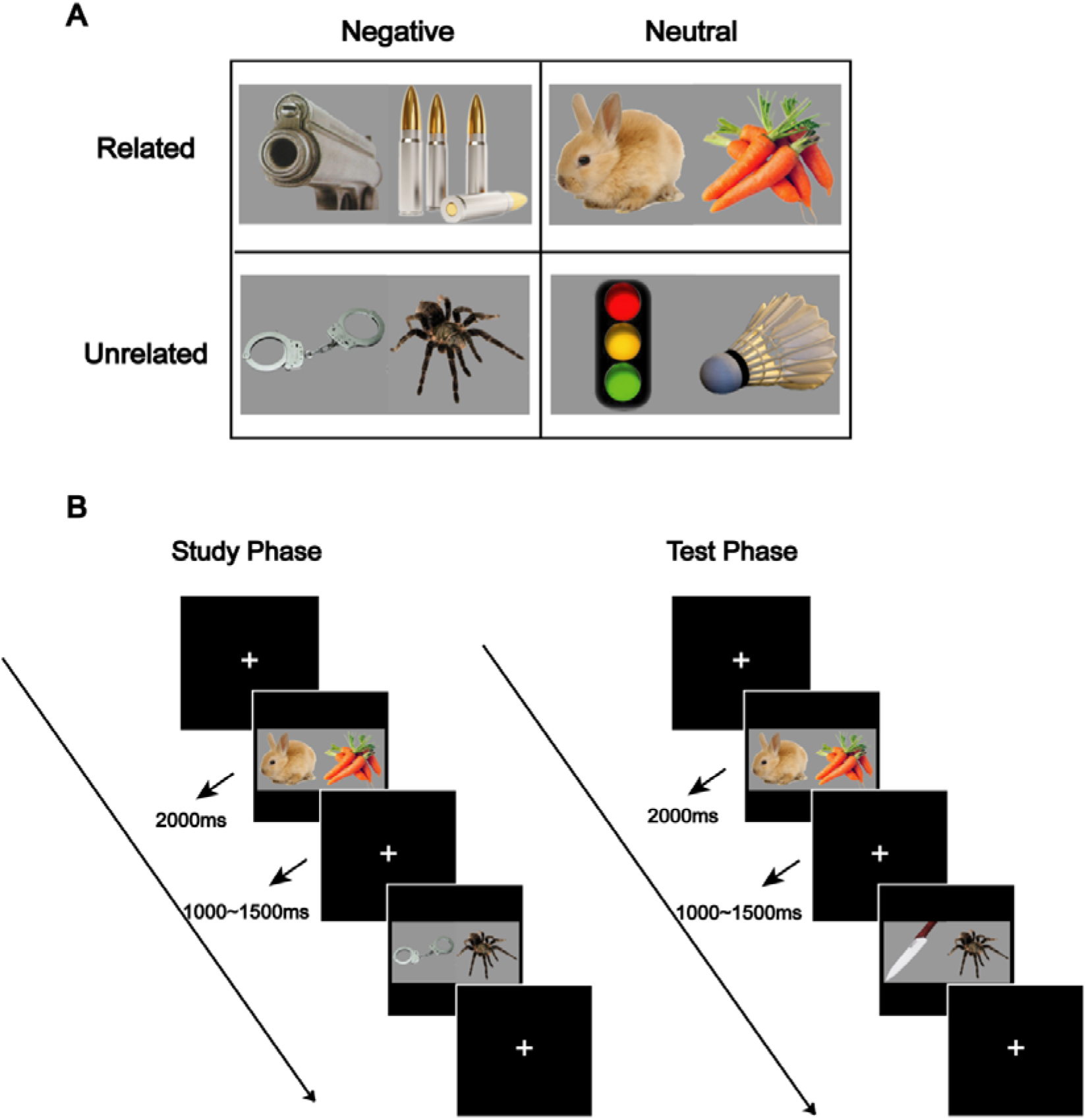
Example stimuli and experimental paradigm. (A) Example stimuli for each condition. Our factorial design included four types of picture pairs, varying in their semantic relatedness (related, unrelated) and valence (negative, neutral). (B) Schematic illustration of the experimental paradigm. At study, participants viewed picture-pairs and indicated which item is more negative (Experiment 1) or more familiar (Experiment 2). At test, they discriminated between intact, rearranged, and new pairs.

An independent sample (N = 12) provided ratings of the pictures on the dimensions of valence, arousal and familiarity, using a scale ranging from 1 (very negative/calm/unfamiliar) to 9 (very positive/exciting/familiar). Pictures with familiarity ratings below 4 were removed from the pool. 1272 pictures, including 646 negative pictures (with valence scores below 4) and 625 neutral pictures (with valence scores between 4 and 7), were chosen and combined to form negative/negative picture pairs or neutral/neutral picture pairs, resulting in 166 semantically-unrelated negative, 150 semantically-unrelated neutral, 157 semantically-related negative, and 162 semantically-related neutral picture pairs. Another independent sample (N = 10) provided ratings of the pairs on the dimension of relatedness. They were asked to judge how likely it is for the two objects to appear together (Tibon et al., 2014), by responding on a scale ranging from 1 (very unlikely) to 9 (very likely). Only pairs for which the pre-assigned relatedness status was verified (i.e., unrelated stimuli that with relatedness score < 5, and related stimuli with relatedness score >= 5) by the majority of the raters (at least 6/10 raters) were included in the study. Based on the ratings, 600 pairs were selected, including 150 semantically-related negative pairs, 150 semantically-related neutral pairs, 150 semantically-unrelated negative pairs, and 150 semantically-unrelated neutral pairs. Semantically-related pairs either belonged to the same category (e.g., “desk-sofa”), or were functionally related (e.g., “rabbit-carrot”). The relatedness scores of related pairs [Mean (SD) = 6.61 (1.25)] were significantly higher than that of unrelated pairs [Mean (SD) = 1.61 (.41); *t* (299) = 64.20, *p* < .001]. Negative pictures were significantly more negative and arousing than neutral pictures [valence: Mean_neg_ (SD) = 3.05 (0.66), Mean_neu_ (SD) = 5.06 (0.31), *t* (599) = -115.45, *p* < .001; arousal: Mean_neg_ (SD) = 5.43 (1.23), Mean_neu_ (SD) = 2.41 (0.63), *t* (599) = 52.36, *p* < .001], but equal to neutral pictures on familiarity (*p* > .05).

We subsequently constructed 600 rearranged pairs by combining together pictures belonging to different pairs, but keeping their type and location unchanged, such that there were 150 rearranged pairs for each type. For example, two related neutral pairs A-B (e.g., rabbit-carrot) and C-D (e.g., goat-cabbage) could be recombined to form another related neutral pair A-D (rabbit-cabbage). B and C would also be combined with other items (belonging to related neutral pairs) to form rearranged pairs. The same sample (N = 10) of participants in prior relatedness ratings were recruited again and provided ratings for relatedness. The results confirmed our initial assignment of pairs, and showed that the relatedness scores of related pairs [Mean (SD) = 6.67 (1.23)] were significantly higher than that of unrelated pairs [Mean (SD) = 1.65 (.57); *t* (299) = 64.90, *p* < .001]. Importantly, there was no difference in relatedness between the rearranged pairs and the original pairs [Unrelated pairs: *t* (299) = 1.07, *p* = 0.29; Related pairs: *t* (299) = 0.76, *p* = 0.35].

A total of 400 picture pairs were encoded at the study phase (100 related negative pairs, 100 related neutral pairs, 100 unrelated negative pairs, 100 unrelated neutral pairs), with the remaining 200 pairs serving as new pairs during the test phase. At test, 200 intact pairs (the same pairs shown at study), 200 rearranged pairs (pictures belonging to different study pairs that were recombined together), and 200 new pairs were presented, with each condition containing 50 related negative, 50 related neutral, 50 unrelated negative, and 50 unrelated neutral pairs. Test pairs were counterbalanced across subjects, with every picture presented equally often as part of intact, rearranged, or new pairing.

#### Procedure

Participants were seated at a distance of 70 cm from a Dell monitor in an electrically shielded room. Picture pairs, with a visual area of 10° × 5°, were displayed (using Presentation by Neurobehavioral Systems, Inc) horizontally at the center of the monitor against a black background. A standard study-test paradigm was adopted, with the study phase followed by the test phase after a 10 min delay. Four self-paced breaks were provided during the study phase and during the test phase. Stimuli were presented pseudo-randomly to ensure that no more than three consecutive trials were from the same condition.

At study, each trial began with a gray fixation cross for 1000∼1500 ms, followed by the presentation of a picture pair for 2000 ms, during which the participants were asked to memorize the pairs and perform a valence judgement task, namely, to judge which one of the two objects is more negative (Fig.1B). They were asked to press the ‘left arrow’ key on the keyboard if they thought that the left object was more negative, to press the ‘right arrow’ key if they thought that the right one was more negative, and to press the ‘down arrow’ key if they thought that the two objects had similar valence. Once the study phase was completed, a 10 min break was provided. During this period, participants performed a distractor task of 3-digit backward counting for 5 min and then rested for five additional minutes.

At test, each trial began with a jittered fixation cross presented for 1000∼1500 ms, followed by the presentation of a picture pair for 2000 ms. Participants were asked to indicate whether the pair is “intact”, “rearranged”, or “new” as accurately and as quickly as possible. Responses were provided via keyboard keys, counterbalanced across participants. Half of the participants made responses of “intact” and ‘‘rearranged” by pressing the key ‘‘F” and ‘‘G” with left hand, and of ‘‘new” by pressing the key ‘‘J” with right hand. The other half of participants responded ‘‘intact” and ‘‘rearranged” by pressing the key ‘‘H” and ‘‘J” with right hand, and ‘‘new” by pressing the key ‘‘F” with left hand.

A study practice block of 12 trials was provided at the beginning of the experiment, prior to the study phase. An additional test practice block of 18 trials was provided prior to the test phase. During these practice sessions, the experimenter ascertained that the participants understood the task.

#### EEG recording and preprocessing

EEG was recorded using a 64-channel Neuroscan system and the electrode locations adhered to the extended international 10–20 system. The sampling rate was 500 Hz with a 0.05∼100 Hz bandpass filter. Electrooculogram (EOG) was recorded using two electrodes placed outside the outer canthi of each eye and one infraorbital to the left eye. The left mastoid was used as the reference site online, and EEG signals were re-referenced offline to the average of the left and right mastoid recordings. Impedance was kept below 5 kΩ. EEG/EOG signals were filtered with a bandpass of 0.05∼40 Hz. EEG data from the test phase were segmented into 1100 ms epochs, corrected to the 100 ms pre-stimulus baseline. Epochs with voltage exceeding ±75 μV were excluded. Independent component analysis (ICA) conducted with the runica algorithm available through the EEGLAB toolbox for MATLAB (v.2019.0, Delorme & Makeig, 2004), was used to isolate and remove EOG blink artifacts. A minimum of 16 trials for each condition was required to ensure an acceptable signal-to-noise ratio, and four participants were excluded for failing to meet the minimal number of trials.

Mean numbers of related analyzed trials were 39 (intact), 26 (rearranged), and 38 (new) for negative pairs, and 37 (intact), 26 (rearranged), and 43 (new) for neutral pairs. Mean numbers of unrelated analyzed trials were 25 (intact), 30 (rearranged), and 37 (new) for negative pairs, and 27 (intact), 30 (rearranged), and 41 (new) for neutral pairs.

#### Statistical analyses: General approach

Data were extracted for correct trials only (e.g., Donaldson & Rugg, 1998; Paller et al., 2003). Repeated measures analyses of variance (ANOVAs) were conducted for inferential statistics, with Greenhouse-Geisser correction for non-sphericity when required. Follow-up analyses were performed using repeated measures ANOVAs or t-tests as appropriate. To control for Type I error rates, p-values were corrected for false discovery rate (FDR) with the Benjamini–Hochberg procedure (Benjamini & Hochberg, 1995) at *p* < .05. Because the current study focuses on mnemonic effects, only main effects and interactions that included the factor of response type are reported.

##### Behavioral analyses

Behavioral data of interest was associative Pr: a discrimination measure of old/new effects for associative memory, defined by subtracting false alarm rates for rearranged pairs from hit rates for intact pairs (Jäger et al., 2006; Snodgrass & Corwin, 1988). This measure was used to dissociate potential response bias (e.g., for related pairs; Ahmad & Hockley, 2014; Liu et al., 2019; Tibon et al., 2014; see Supp. Mat. 1 for an ancillary analysis of response bias) from a true memory advantage. Pr scores were analyzed using a repeated measure ANOVA with relatedness (related or unrelated) and valence (negative or neutral) as repeated factors, and with Pr score as the dependent measure.

Given our interest in associative memory, and to disentangle true memory effects from response bias, we further analyzed accuracy rates (% correct) for intact pairs and for rearranged pairs using a repeated ANOVA with relatedness (related or unrelated) and valence (negative or neutral) as repeated factors, and accuracy rate as the dependent measure. A full 3-way ANOVA which includes all factors (relatedness, valence, and response type) and levels (intact, rearranged, new) within the same model, is included in the supplementary materials (Supp. Mat. 2).

##### ERP analyses

Both intact and rearranged pairs are comprised of studied items. However, while intact pairs further contain studied associative information, rearranged pairs contain novel associative information which was not presented at study. Therefore, the intact/rearranged effects, i.e., differences between ERPs associated with correct “intact” judgments vs. correct “rearranged” judgments, is indicative of associative recognition. (e.g., Li et al., 2017; Rhodes & Donaldson, 2008; Zheng et al., 2015). Therefore, we focused on the comparison between intact and rearranged pairs to index associative memory. For completion, we also include the comparison between rearranged and new pairs as an index of item memory in the supplementary materials (Supp. Mat. 3).

For the frontal and parietal memory effects, time segments and regions of interest were defined based on previous ERP studies (Bader et al., 2010; Han et al., 2018; Li et al., 2017, 2019; Rugg & Curran, 2007; Speer & Curran, 2007; Wolk et al., 2006; Zheng et al., 2016). Accordingly, two time windows, 300-550 ms and 550-800 ms, were used to capture the frontal and parietal memory effects, respectively. Mean amplitudes for statistical analyses in these windows were obtained from frontal (collapsed across F3, Fz, and F4), central (collapsed across C3, Cz, and C4) and parietal (collapsed across P3, Pz, and P4) scalp locations (Han et al., 2018; Hou et al., 2013; Molinaro et al., 2011).

Repeated measures ANOVA was conducted separately for each time window and included four within-subjects factors: relatedness (related or unrelated), valence (negative or neutral), response type (intact or rearranged), and location (frontal, central, or parietal).

### Results

#### Behavioral results

Means and SDs for the various behavioral measures of Experiment 1 are shown in Table 1. The ANOVA for associative Pr (relatedness × valence) revealed a significant main effect of valence, *F* (1, 41) = 24.28, *p* < .001, η^2^_p_ = 0.37 (greater Pr scores for neutral vs. negative pairs), and of relatedness, *F* (1, 41) = 120.01, *p* < .001, η^2^_p_ = 0.75 (greater Pr scores for related vs. unrelated pairs). The analysis further revealed a significant 2-way interaction between relatedness and valence, *F* (1, 41) = 6.61, *p* = .014, η^2^_p_ = 0.14, with lower associative Pr for negative pairs (vs. neutral) in the unrelated condition, *t* (41) = 5.38, *p* < .001, d = 0.83, but not in the related condition, *t* (41) = 1.65, *p* = 0.107, d = 0.25.

**Table 1.**
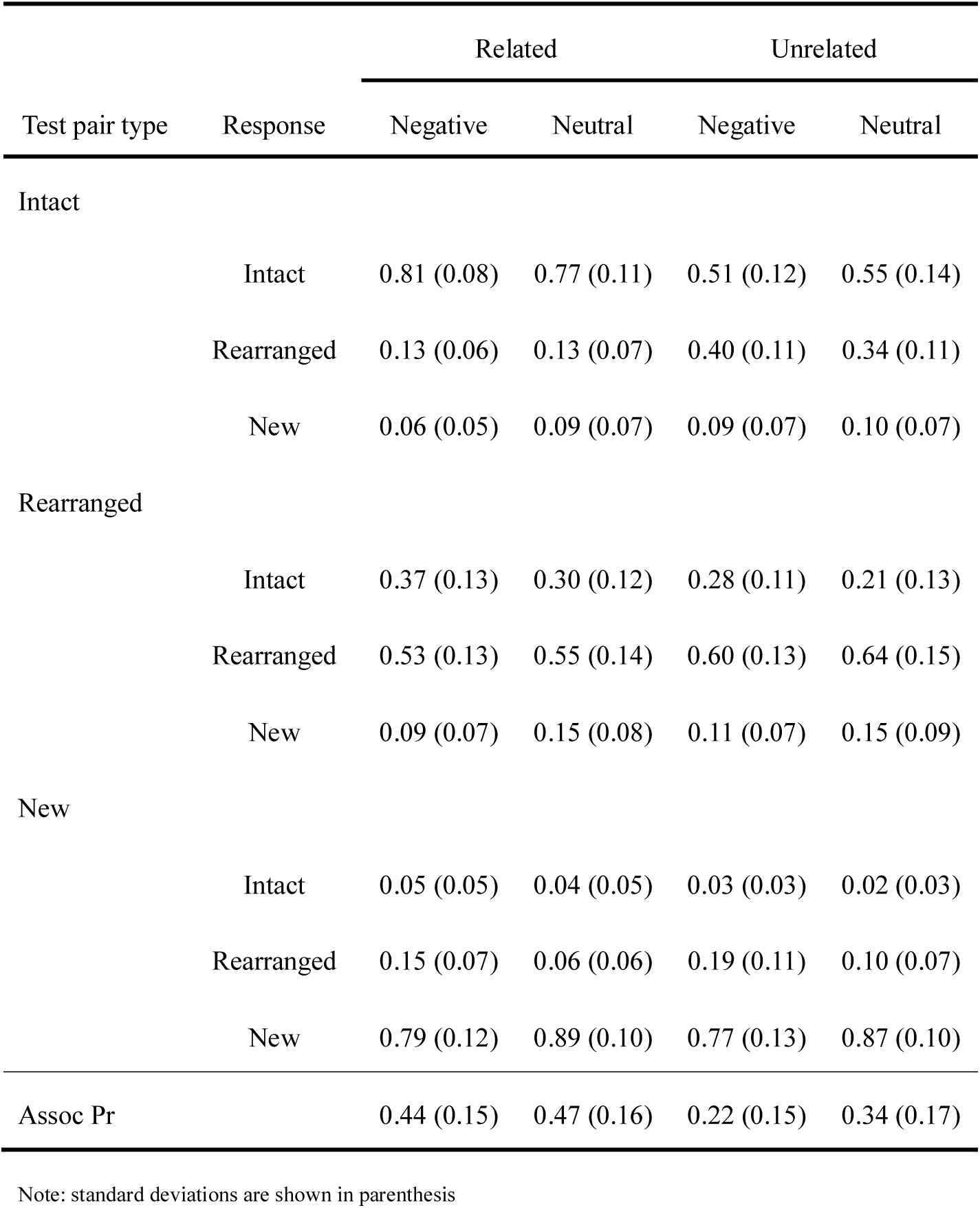
Distribution of participants’ responses and Pr scores in Experiment 1

The analysis of accuracy rates for intact pairs (“hits”) revealed a main effect of relatedness, *F* (1, 41) = 361.91, *p* < .001, η^2^_p_ = 0.90, and a 2-way interaction between the two factors, *F* (1, 41) = 17.35, *p* < .001, η^2^_p_ = 0.30, resulting from lower accuracy rates for negative pairs (vs. neutral) in the unrelated condition, *t* (41) = 2.78, *p* = .008, d = 0.43, but greater accuracy rates for negative pairs (vs. neutral) in the related condition, *t* (41) = 2.31, *p* = .026, d = 0.36. The analysis of accuracy rates for rearranged pairs (“correct rejections") only revealed a main effect of relatedness, *F* (1, 41) = 22.70, *p* < .001, η^2^_p_ = 0.36, with greater accuracy rates for unrelated vs. related pairs.

Taken together, the behavioral results depict the predicted emotional associative interference effect, indicated by reduced Pr scores and accuracy rates for negative vs. neutral pairs. Furthermore, this effect was attenuated by semantic relatedness, with greater emotional interference observed for unrelated vs. related pairs.

#### ERP results

Waveforms and topographical distribution of the associative memory effect in the various experimental conditions are shown in Figure 2. In the early time window (300-550 ms), the ANOVA for the associative memory effect revealed a 2-way interaction between valence and response type, *F* (1, 41) = 9.29, *p* = .004, *η^2^_p_* = 0.19. Decomposition of the interaction using paired t-tests at each level of valence, revealed a significant associative memory effect (more positive-going waveforms for intact vs. rearranged) for negative pairs, *t* (41) = 3.13, *p* = .003, d = 0.48, but not for neutral pairs (*p* = 0.29). Thus, in the early time-window, the associative memory effect emerged for negative pairs, regardless their relatedness, and had a widespread distribution. In the late time window (550-800 ms), the ANOVA only revealed a main effect of response type, *F* (1, 41) = 38.84, *p* < .001, ^2^ = 0.49, suggesting that the late associative memory effect was similarly observed in all conditions, and in all locations. Exploratory analysis of a later associative memory effect (800-1,000 ms), which resembled the pattern observed in the 550-800 ms time window, is included in the supplementary materials (Supp. Mat. 4).

**Figure 2.**
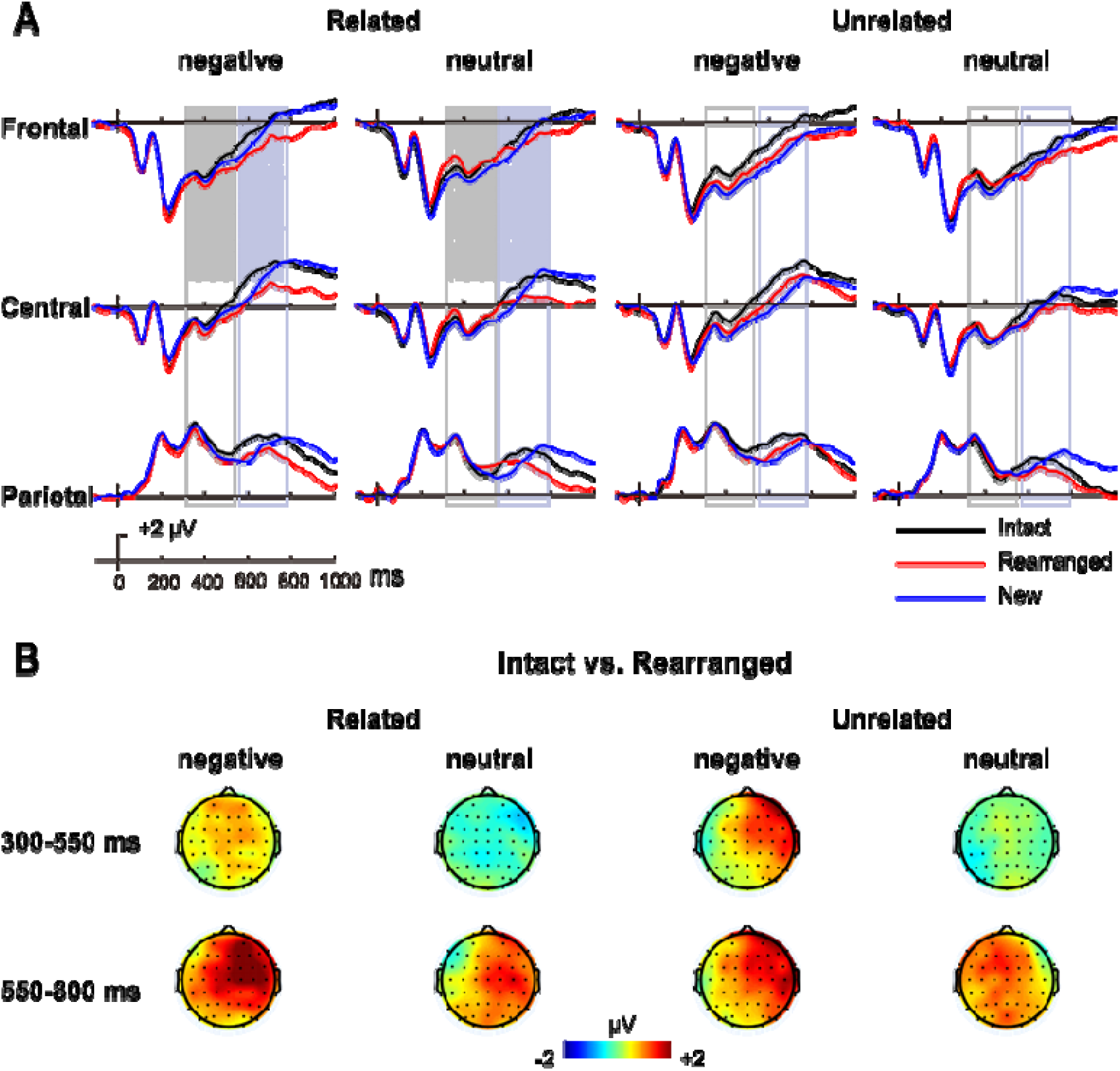
ERP results for Experiment 1. (A) Grand average ERP waveforms for intact responses (black), rearranged responses (red), and new responses (blue) in the four experimental conditions (related\unrelated × negative\neutral) at three scalp locations (F3, Fz, F4 collapsed as the frontal site; C3, Cz, C4 as the central site; P3, Pz, P4 as the parietal site). Shaded areas indicate time-windows used for the analyses of the early memory effects (300-550 ms, light grey) and the late memory effects (550-800 ms, dark gray). (B) Topographical maps of the associative memory effects (intact minus rearranged) in each time window.

### Discussion

In Experiment 1 we observed an associative emotional interference effect, which was attenuated by semantic relatedness. Turning to the EEG data, semantic relatedness did not modulate the early associative memory effect, associated with familiarity, nor the late associative memory effect, associated with recollection. That is, the difference waveform for intact vs. rearranged pairs was similar for related and unrelated pairings. Interestingly, however, the associative memory ERP effect was modulated by the valence of the stimuli: the early ERP component showed a frontal modulation for negative pairs (with greater frontal negativity for rearranged vs. intact pairings), suggesting unitizability of negative stimuli and the contribution of familiarity to their associative recognition. Nevertheless, even though the modulation of the early ERP effect was similar for related and unrelated pairs, the behavioral attenuation of the associative emotional interference effect was only apparent for semantically related. We return to this seemingly discrepancy between the behavioral and the ERP results in our General Discussion.

Although somewhat unpredicted, the modulation of the early ERP component for negative (but not neutral) pairs, adheres to the suggestion that valence serves as an organizing principle, i.e., allowing grouping across shared properties. Previous studies (Talmi & Moscovitch, 2004; Talmi et al., 2007) proposed that the common finding of enhanced memory for emotional items can be due to a shared emotional context. In a series of experiments, they showed that the memory advantage for emotional items is eliminated when these are compared with categorized neutral items, suggesting that valence serves as an organizing principle for the items. This organization, along the dimension of emotional valence, possibly overshadowed the effect of semantic relatedness.

Indeed, in the current experiment, emotional content was explicitly probed (i.e., during study participants completed valence judgments for the stimuli), which might have promoted emotional processing and hindered processing of semantic relations. Notably, a growing number of evidence suggests that modulation of behavior or processing by emotional content is only triggered when emotions are relevant to the task (e.g., Engen et al., 2017; Huang et al., 2008; Pessoa, 2009; Stein et al., 2009; Wei et al., 2015). For example, Stein et al (2009) conducted the attentional blink task to estimate whether task relevance impacts prioritization. In this experiment, participants judged either the emotion (relevant condition) or gender (irrelevant condition) of two target facial stimuli depicting different expressions (fearful or neutral). Fearful faces (vs. neutral) induced a stronger attentional blink, but only in the relevant condition. This demonstrates that the processing advantage of emotional stimuli depends on the relevance of emotion to the task. Similarly, in the context of the current task, situating emotion as task-relevant might have promoted binding along this dimension. In other words, instead of creating unitized links for semantically related (vs. unrelated) items, such links were created for negative (vs. neutral) ones. Experiment 2 was designed to address this possibility.

## Experiment 2

In Experiment 2, we investigated the effect of semantic relatedness on associative recognition of emotional stimuli, when emotions are incidental to the task. To this end, Experiment 2 employed the same behavioral and EEG measures as in Experiment 1, but with a different study task. Namely, rather than valence judgment, in Experiment 2 participants were asked to perform a familiarity judgment.

We predicted an overall behavioral pattern similar to Experiment 1, that is, a smaller associative emotional interference effect for semantically related vs. unrelated pairs. We further rationalized that, with emotion being incidental to the task, both unitization and LOP would be promoted for semantically related (vs. unrelated) pairs. We therefore predicted that both the early and the late associative memory ERP effects would be greater for related than unrelated pairs.

### Method

#### Participants

Forty-seven healthy, right-handed native Chinese speakers (32 females; mean age 22.1 ± 1.9 years) from Capital Normal University participated, with the same characteristics as those who participated in Experiment 1, participated in Experiment 2. Data from seven participants were discarded, due to insufficient number of artifact-free ERP trials in one or more experimental conditions (N < 16). Our final sample therefore included 40 participants (27 females; mean age 21.9 ± 2.0 years).

#### Procedure, recording, and analyses

Stimuli were the same as in Experiment 1. The procedure was identical to that of Experiment 1, except that participants were asked to perform a familiarity judgement task at study, namely, to judge which one of the two objects presented in each trial was more familiar. They were asked to press the ‘left arrow’ key on the keyboard if they thought that the left object was more familiar, to press the ‘right arrow’ key if they thought that the right one was more familiar, and to press the ‘down arrow’ key if they thought that the two objects do not differ in their familiarity.

EEG recording and preprocessing were the same as in Experiment 1. Mean numbers of related analyzed trials were 38 (intact), 26 (rearranged), and 39 (new) for negative pairs, and 40 (intact), 29 (rearranged), and 43 (new) for neutral pairs. Mean numbers of unrelated analyzed trials were 23 (intact), 29 (rearranged), and 36 (new) for negative pairs, and 28 (intact), 31 (rearranged), and 42 (new) for neutral pairs. Statistical analyses were the same as in Experiment 1.

### Results

#### Behavioral results

Means and SDs for the various behavioral measures of Experiment 2 are shown in Table 2. The ANOVA for associative Pr revealed main effects of valence, *F* (1, 39) = 40.31, *p* < .001, η^2^_p_ = 0.51, and relatedness, *F* (1, 39) = 220.81, *p* < .001, η^2^_p_ = 0.85, with greater Pr scores for neutral pairs (vs. negative), and for related pairs (vs. unrelated).

**Table 2.** Distribution of participants’ responses and Pr scores in Experiment 2

| Test pair type | Response | Related |  | Unrelated |  |
| --- | --- | --- | --- | --- | --- |
|  |  | Negative | Neutral | Negative | Neutral |
| Intact |  |  |  |  |  |
|  | Intact | 0.79 (0.08) | 0.83 (0.10) | 0.48 (0.12) | 0.59 (0.14) |
|  | Rearranged | 0.16 (0.06) | 0.11 (0.05) | 0.43 (0.10) | 0.34 (0.10) |
|  | New | 0.06 (0.05) | 0.06 (0.07) | 0.09 (0.07) | 0.07 (0.07) |
| Rearranged |  |  |  |  |  |
|  | Intact | 0.36 (0.12) | 0.29 (0.11) | 0.26 (0.09) | 0.27 (0.12) |
|  | Rearranged | 0.54 (0.12) | 0.60 (0.13) | 0.62 (0.12) | 0.64 (0.14) |
|  | New | 0.10 (0.05) | 0.11 (0.09) | 0.12 (0.07) | 0.09 (0.06) |
| New |  |  |  |  |  |
|  | Intact | 0.05 (0.06) | 0.02 (0.04) | 0.04 (0.04) | 0.01 (0.02) |
|  | Rearranged | 0.15 (0.08) | 0.07 (0.05) | 0.21 (0.12) | 0.11 (0.08) |
|  | New | 0.80 (0.12) | 0.90 (0.08) | 0.74 (0.14) | 0.89 (0.10) |
| Assoc Pr |  | 0.43 (0.13) | 0.54 (0.15) | 0.22 (0.12) | 0.31 (0.18) |

Note: standard deviations are shown in parenthesis

The analysis of accuracy rates for intact pairs revealed a main effects of relatedness, *F* (1, 39) = 370.64, *p* < .001, η^2^_p_ = 0.91, and valence, *F* (1, 39) = 22.64, *p* < .001, η^2^_p_ = 0.37, as well as a 2-way interaction between the two factors, *F* (1, 39) = 12.67, *p* = .001, η^2^_p_ = 0.25. Decomposition of this interaction showed that even though the difference in accuracy rates between neutral and negative pairs emerged for both related and unrelated pairs, it was greater in the latter, *t* _related_(39) = 2.35, *p* = .024, d = 0.37; *t _un_*_related_(39) = 5.26, *p* < .001, d = 0.83. For rearranged pairs, the analysis of accuracy rates revealed a main effect of relatedness, *F* (1, 39) = 12.88, *p* = .001, η^2^_p_ = 0.25, and valence, *F* (1, 39) = 6.59, *p* = .014, η^2^_p_ = 0.15, but no significant interaction.

The behavioral results of Experiment 2 show a similar pattern of attenuated emotional interference effect for related vs. unrelated pairs, which we observed in Experiment 1. However, unlike Experiment 1, in which this pattern was observed both for hit rates (correct “intact” responses) and for the unbiased associative Pr scores, in Experiment 2 it was only observed for the former.

#### ERP results

Waveforms and topographical distribution of the associative memory effect are shown in Figure 3. In the early time window (300-550 ms), the analysis revealed a main effect of response type, with more positive-going waveforms for intact pairs (vs. rearranged), *F* (1, 39) = 4.67, *p* = .037, η^2^_p_ = 0.11, and a 2-way interaction between relatedness and response type, *F* (1, 39) = 5.09, *p* = .030, η^2^_p_ = 0.12. Decomposition of the interaction showed a significant associative memory effect for related pairs, *t* (39) = 3.08, *p* = .004, d = 0.49, but not for unrelated pairs (*p* = 0.81). Thus, in the early time-window, the associative memory effect emerged for related pairs, regardless their valence, and had a widespread distribution.

**Figure 3.**
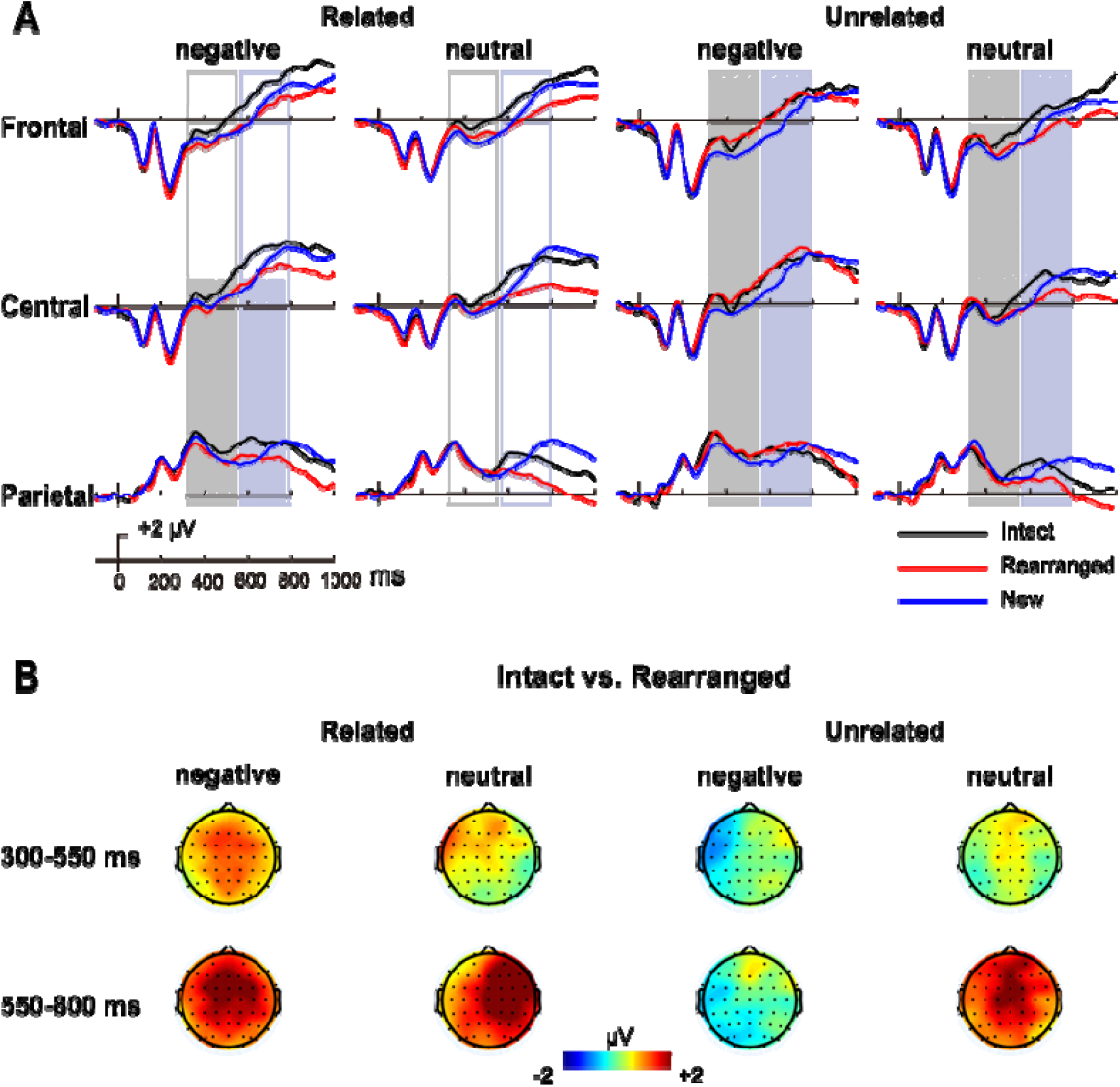
ERP results for Experiment 2. (A) Grand average ERP waveforms for intact responses (black), rearranged responses (red), and new responses (blue) in the four conditions at three scalp locations. Shaded areas indicate time-windows used for the analysis of the early and late memory effects. (B) Topographical maps of the associative memory effects in each time window.

In the late time window (550-800 ms), the analysis revealed a main effect of response type, *F* (1, 39) = 38.49, *p* < .001, *η^2^_p_* = 0.50, a 2-way interaction between relatedness and response type, *F* (1, 39) = 10.23, *p* = .003, *η^2^_p_* = 0.21, a 2-way interaction between valence and response type, *F* (1, 39) = 6.92, *p* = .012, *η^2^_p_* = 0.15, and a 3-way interaction between relatedness, valence, and response type, *F* (1, 39) = 8.37, *p* = .006, *η^2^_p_* = 0.18. To decompose the 3-way interaction, we conducted follow-up ANOVAs with relatedness and response type as within-subject factors, separately for each valence. For negative pairs, this analysis revealed a main effect of response type, *F* (1, 39) = 11.46, *p* = .002, *η^2^_p_* = 0.23, and a significant 2-way interaction between relatedness and response type, *F* (1, 39) = 17.12, *p* < .001, *η^2^_p_* = 0.31, resulting from an associative memory effect for related pairs, *t* (39) = 4.74, *p* < .001, d = 0.75, but not for unrelated pairs (*p* = .82). A similar follow-up ANOVA for neutral pairs revealed a main effect for response type, *F* (1, 39) = 42.93, *p* < .001, *η^2^_p_* = 0.52, but no interaction between the factors. As for Experiment 1, an exploratory analysis of a later associative memory effect (800-1,000 ms) is included in the supplementary materials (Supp. Mat. 4).

### Discussion

In Experiment 2, participants were more likely to classify negative pairs as intact if they were semantically related, but this similarly occurred for both intact and rearranged pairs. We come back to this in the General Discussion below.

As for the ERP data, the results agree with previous studies (e.g., Ahmad & Hockley, 2014; Kriukova et al., 2013; Li et al., 2019; Rhodes & Donaldson, 2008; Tibon et al., 2014), demonstrating modulation of the early associative memory effect by semantic relatedness. Namely, the early difference between intact and rearranged pairs was only significant when stimuli were semantically related (regardless their valence). The late associative memory effect was more generally distributed, and was only absent for unrelated negative pairs. While this absence might seem puzzling at first sight, a closer look at the behavioral results suggests that this is the only experimental condition (across both experiments) for which the probability to correctly classify intact items (47%) was highly similar to the probability to classify them as rearranged pairs (43%). This suggests that in this condition, for which emotional associative interference is not attenuated by semantic relations, recollective processes are impeded. Yet, it remains unclear why the same pattern was not observed in Experiment 1.

### General Discussion

The present study used an associative recognition paradigm, across two experiments, to investigate how semantic relations, valence, and their interaction, affect our ability to retrieve associative information. Both experiments conformed to the same procedure, aside from the instructions given during the study phase. Namely, in Experiment 1 participants were asked to compare the valence of object pairs, whereas in Experiment 2 they were asked to compare their familiarity.

In both experiments, an associative emotional interference effect had emerged, with reduced associative memory for negative pairs. This finding agrees with previous studies, showing that emotion can impair associative memory (Madan et al., 2012; Mao et al., 2015; Mather & Knight, 2008; Pierce & Kensinger, 2011; Rimmele et al., 2011). Importantly, our results further suggest that this associative emotional interference effect can be reduced under certain circumstances. In particular, when valence is attended (as in the case of Experiment 1, where the task requires valence judgment) semantic associative relations can attenuate this associative emotional interference effect. Interestingly, in Experiment 2, negative related pairs were more likely to be classified as “intact” compared to negative unrelated pairs (as indicated by their accuracy scores). This tendency, however, was not reflected in the unbiased associative Pr scores. Therefore, in the case of Experiment 2, the alleviated number of “intact” responses does not represent a memory effect, but rather indicates a response bias. Taken together, the results of the two experiments agree with previous research, showing that emotions can enhance processing, but only when they are relevant to the task (as in Experiment 1; e.g., Engen et al., 2017; Huang et al., 2008; Pessoa, 2009; Stein et al., 2009; Wei et al., 2015). In the current case, during encoding, processing of semantic relations was enhanced for negative pairs, leading to better binding which reduced the associative emotional interference. Nevertheless, this reduction only occurred when emotions were task-relevant (i.e., in Experiment 1).

Turning to the ERP data, our results revealed that the modulation of the early associative memory effect was task dependent. Specifically, in Experiment 1, where valence was relevant to encoding, the early ERP effect showed greater frontal negativity for intact vs. rearranged negative pairs, regardless their semantic relations. In contrast, in Experiment 2, where familiarity (and not valence) was probed during encoding, this modulation only occurred for related (but not for unrelated) pairs, regardless their valence. These results provide evidence for the suggestion that emotions triggered by stimulus’ valence can serve as an organizing principle that binds the items together via shared context (Riberto et al., 2019; Talmi & Moscovitz, 2004). Therefore, pairs of emotional stimuli might be more easily unitized compared to non-emotional ones. The results of Experiment 1 support this idea. Namely, in this experiment, the early associative memory effect—the putative electrophysiological correlate of familiarity—showed a modulation for negative pairs, indicating that familiarity was readily available for these pairs.

Interestingly, this valence-based modulation of the early effect was not observed in Experiment 2, in which valence was not probed during encoding, suggesting that valence might only serve as an organizing principle when it is task relevant (or attended). When valence is not relevant to the task, familiarity signals can be elicited when the stimuli comprising the pair are semantically related, as apparent in Experiment 2. Indeed, previous studies have shown that semantic relations can promote unitization, which further enhances familiarity-based associative recognition (e.g., Ahmad & Hockley, 2014; Kriukova el al., 2013; Li et al., 2019; Rhodes & Donaldson, 2008; Tibon et al., 2014). Arguably, when valence is irrelevant, semantic relatedness “pops-out” as an organizing principle, and enables unitization along this dimension. Taken together, the modulation of the early associative effect observed in the current study, suggests that both semantic relationships and emotional context can support unitization which can subsequently promote familiarity-based retrieval.

Unlike the selective modulation of the early associative memory effects, the modulation of the late effect was apparent more generally across the various experimental conditions (albeit, as noted above, not for unrelated negative pairs in Experiment 2). Although this later effect was rather broadly distributed and lacked the pronounced parietal maxima often associated with the recollection-related late positive component (Wilding & Ranganath, 2011; Rugg & Curran, 2007), retrieval-related modulations with anterior/central topographic distribution are commonly reported in associative recognition ERP studies (e.g., Bader et al., 2010; Han et al., 2018; Kriukova et al., 2013; Mollison & Curran, 2012; Rhodes & Donaldson, 2007, 2008; Tibon, Ben-Zvi, & Levy, 2014; Zheng, Li, Xiao, Broster, & Jiang, 2015) and are interpreted as reflecting recollective processes. Our findings suggest that recollective processes were readily available for intact pairs, regardless their valence or semantic relations.

One aspect of the associative memory effects that warrants further attention is the correspondence between the behavioral results and the ERP data. Specifically, in Experiment 1, associative emotional interference was only attenuated when semantically related pairs were retrieved, even though the modulation of the early ERP effect was similar for related and unrelated pairs. We speculate that the production of early mnemonic signals (as indicated in the ERPs) would only affect behavior if these signals are considered diagnostic or trusted. For semantically related pairs, the emotional context adjoins the semantic one (e.g., for gun:bullets pair: negative feeling due to the shooting gun), producing a trustworthy mnemonic signal that attenuates associative emotional interference. In contrast, for stimuli that lack semantic relations, the early signal is cognitively attributed to more general sources (e.g., that the stimuli were experienced together during the experimental session, rather than within specific pairing). Therefore, these signals are not designated as trustworthy, and do not produce the same behavioral change. This idea coincide with a recent proposal by Bastin et al (2019) which posits that an attribution system modulates the use of memory traces as a function of expectancies, task context, and goals, leading to subjective experiences and explicit judgments. In the current case, associative familiarity signals might have been attributed to an alternative source (e.g., such as processing fluency or global familiarity) and were discarded in the face of explicit judgments.

In the current study, the distinction between the two processes supporting recognition memory—familiarity and recollection—relies mainly on the temporal distribution, and to some extent also on the spatial distribution, of the early and late intact/rearranged ERP effects. Like many other studies (e.g., Bader et al., 2010; Guillaume & Etienne, 2015; Jäger et al., 2006; Kamp et al., 2016; Rhodes & Donaldson, 2008; Tibon et al., 2014; Zheng et al., 2015, to list just a few), we associate the early frontal negativity with familiarity, and the late positivity with recollection. While this type of reverse inference has its limitations (Poldrack, 2006), our interpretation builds on decades of intensive research that strongly associates these ERP components with the particular memory processes reported here (reviewed by Mecklinger, 2000; Rugg & Curran, 2007; Wilding & Ranganath, 2011). We do acknowledge that in the current study, the early ERP component might reflect processing fluency instead of (or possibly together with) familiarity (see Paller et al., 2007; Paller et al., 2012; Mecklinger et al., 2012 for discussion). In any event, however, even if the links made in our study between electrophysiological measures and specific recognition processes are not entirely conclusive (though strongly suggestive), our data clearly point to a neural distinction, whereby the contribution of the early ERP effect to associative recognition is limited to certain experimental conditions, but the contribution of the late ERP effect is more widely available across different types of associated information.

One caveat of the current study, is that negative stimuli were also highly arousing (compared to neutral, see Stimuli section above). Therefore, one potential difficulty in interpretation of the present findings is that valence effects cannot be distinguished from arousal effects, even though prior studies suggest that these might rely on distinct neural process (e.g., Kensinger & Corkin, 2004). Furthermore, the current study only included negative and neutral stimuli, precluding any conclusions regarding general emotional effects, or distinction between different kinds of valence. In addition, the study did not include any additional emotional measures, such as mood assessment or assessment of anxiety/stress symptoms, to be used as covariates in statistical analyses. Future studies are thus required in order to generalize our conclusions further.

In summary, the current study shows that when items share a context during their encoding, be that semantic or emotional, their associative retrieval can provoke an early neural modulation, arguably indicative of familiarity-based retrieval, which accompanies recollection. This modulation further depends on the way information was encoded: when valence is relevant to encoding, it acts as an organizing principle that triggers early retrieval processes. But when valence is not relevant, other relations (in our case, semantic) serve to organize information. We propose that the neural modulation only relates to behavioral change when the signals are being interpreted as trustworthy, particularly, when pre-existing semantic relations between the various pieces of information are present. This suggests that in real-life situations, where emotional information is often highly relevant and semantically meaningful, activation of early mnemonic signals can be tightly linked to memory performance.

## Author notes

The present study was supported by the National Natural Science Foundation of China (31671127), Support by Capacity Building for Sci-Tech Innovation – Fundamental Scientific Research Funds (No. 025185305000/200). RT was supported by a British Academy Postdoctoral Fellowship (grant SUAI/028 RG94188). We thank Deborah Talmi and Zara Bergström for insightful discussion. The authors declare no conflict of interest.

## Open Practices Statement

The studies reported in this manuscript were not preregistered. Materials for the experiments reported here are available on OSF (https://osf.io/7kpq6/). Data will be available upon request via the MRC Cognition and Brain Sciences Unit data repository.

