## Supplementary Materials for "Effects of emotion and semantic relatedness on recognition memory: Behavioral and electrophysiological evidence"

(1) **Response bias**

Our study shows that in both experiments, negative pairs were more likely to be identified as intact if they were semantically related. Nevertheless, in Experiment 2 this effect was similar for rearranged and intact pairs, whereas in Experiment 1 this effect was greater for intact pairs. This was taken to suggest that a genuine memory effect in Experiment 1, but not in Experiment 2, and that the increased number of “intact” responses in Experiment 2 is therefore driven by response bias. As an additional step, we sought to investigate whether response bias differs across the experimental conditions, and whether our conditions for interest differ in their response bias across the two experiments. We defined response bias as:

Response bias = false alarm rates / (1- Pr)

where ‘false alarm rates’ refer to the percentage of rearranged pairs that were misjudged as intact. For Experiment 1, a repeated measures ANOVA on these scores, with relatedness and valence as repeated factors, revealed a significant main effects of valence, *F* (1, 39) = 12.77, *p* = .001, η^2^p = 0.24, and relatedness, *F* (1, 39) = 186.01, *p* < .001, η^2^p = 0.82, with greater response bias for negative vs. neutral pairs, and for related vs. unrelated pairs. For Experiment 2, the ANOVA revealed a significant main effect of relatedness, *F* (1, 39) = 160.19, *p* < .001, η^2^p = 0.80 (greater for related vs. unrelated pairs), and a 2-way interaction between relatedness and valence, *F* (1, 39) = 7.54, *p* = .009, η^2^p = 0.16. Decomposition of this interaction showed that even though the difference in response bias between related and unrelated pairs emerged for both negative and neutral pairs, it was greater in the former, *t* _negative_(39) = 13.44, *p* < .001, d = 2.13; *t* _neutral_(39) = 8.81, *p* < .001, d = 1.39.

Supp. Table 1. Response bias in Experiments 1 and 2

|  | Related | | Unrelated | |
| --- | --- | --- | --- | --- |
|  | Negative | Neutral | Negative | Neutral |
| Experiment 1 | 0.66 (0.14) | 0.57 (0.18) | 0.36 (0.12) | 0.31 (0.17) |
| Experiment 2 | 0.62 (0.14) | 0.62 (0.16) | 0.33 (0.10) | 0.40 (0.15) |

Note: standard deviations are shown in parenthesis

Next, we computed a difference score for negative pairs by subtracting response bias scores for unrelated negative pairs from that of related negative pairs. After computing this measure, we used an independent sample t-test, to test whether the two experiments differ in the magnitude of response bias for negative related vs. unrelated pairs. This analysis did not reveal a significant difference between the two experiment, *t*(80) = 0.02, *p* = 0.988, d’ = 0.003, with similar response bias for negative related vs. unrelated pairs in Experiment 2 and in Experiment 1. Note, however, that an experimental condition might modulate *both* memory *and* bias. Thus, even though in both studies response bias for negative related pairs was greater than that of negative unrelated pairs (as the ANOVA analyses revealed), the emotional interference memory effect was only reduced in Experiment 1.

(2) **Analyses of accuracy rates**

In addition to the analysis of hit rates (for intact pairs) and correct rejection rates (for rearranged pairs), presented in the main text, we analyzed accuracy rates (% correct) for all response types within the same model. To this end, we used a repeated measured ANOVA with relatedness (related or unrelated), valence (negative or neutral) and response type (intact, rearranged, or new) as repeated factors.

**Experiment 1**

The ANOVA (relatedness × valence × response type) on accuracy rates revealed main effects of relatedness, *F* (1, 41) = 148.19, *p* < .001, *η*^2^_p_ = 0.78, valence, *F* (1, 41) = 34.65, *p* < .001, *η*^2^_p_ = 0.46, with greater accuracy rates for related pairs (vs. unrelated), and for neutral pairs (vs. negative). A main effect also emerged for response type, *F* (2, 82) = 104.41, *p* < .001, *η*^2^_p_ = 0.72, with greater accuracy rates for new pairs vs. intact, *t* (41) = 10.94, *p* < .001, d = 1.69, for intact pairs vs. rearranged, *t* (41) = 4.22, *p* < .001, d = 2.14, and for new pairs vs. rearranged: *t* (41) = 13.87, *p* < .001, d = 0.65. There was also a 2-way interaction between relatedness and response type, *F* (1.63, 66.69) = 140.70, *p* < .001, *η*^2^_p_ = 0.77, a 2-way interaction between valence and response type, *F* (2, 82) = 13.46, *p* = < .001, *η*^2^_p_ = 0.25, a 2-way interaction between relatedness and valence, *F* (1, 41) = 9.26, *p* = .004, *η*^2^_p_ = 0.18, as well as a 3-way interaction between all three factors, *F* (2, 82) = 6.68, *p* = .002, *η*^2^_p_ = 0.14.

We decomposed the 3-way interaction using separate follow-up ANOVAs for each response type, with the factors of relatedness and valence. For intact pairs, this analysis revealed a main effect of relatedness, *F* (1, 41) = 361.91, *p* < .001, *η*^2^_p_ = 0.90, and a 2-way interaction between the two factors, *F* (1, 41) = 17.35, *p* < .001, *η*^2^_p_ = 0.30, resulting from lower accuracy rates for negative pairs (vs. neutral) in the unrelated condition, *t* (41) = 2.78, *p* = .008, d = 0.43, but greater accuracy rates for negative pairs (vs. neutral) in the related condition, *t* (41) = 2.31, *p* = .026, d = 0.36. For rearranged pairs, this analysis revealed a main effect of relatedness, *F* (1, 41) = 22.70, *p* < .001, *η*^2^_p_ = 0.36. For new pairs this analysis revealed main effects of relatedness, *F* (1, 41) = 7.45, *p* = .009, *η*^2^_p_ = 0.15, and valence, *F* (1, 41) = 77.81, *p* < .001, *η*^2^_p_ = 0.66.

**Experiment 2**

For accuracy rates, the analysis revealed main effects of relatedness, *F* (1, 39) = 198.92, *p* < .001, *η*^2^_p_ = 0.84, and valence, *F* (1, 39) = 97.18, *p* < .001, *η*^2^_p_ = 0.71, with greater accuracy rates for related pairs (vs. unrelated), and for neutral pairs (vs. negative). A main effect was also revealed for response type, *F* (2, 78) = 111.76, *p* < .001, *η*^2^_p_ = 0.74, with greater accuracy rates for new vs. intact pairs, *t* (39) = 10.69, *p* < .001, d = 1.69, and vs. rearranged pairs, *t* (39) = 13.82, *p* < .001, d = 2.19, and for intact vs. rearranged pairs, *t* (39) = 4.47, *p* < .001, d = 0.70. There were also 2-way interactions between relatedness and response type, *F* (1.51, 59.00) = 151.17, *p* < .001, *η*^2^_p_ = 0.80, between valence and response type, *F* (2, 78) = 9.52, *p* < .001, *η*^2^_p_ = 0.20, and between relatedness and valence, *F* (1, 39) = 6.84, *p* = .013, *η*^2^_p_ = 0.15. Finally, the analysis revealed a significant 3-way interaction between all 3 factors, *F* (2, 78) = 8.52, *p* < .001, *η*^2^_p_ = 0.18.

Decomposition of the 3-way interaction, separately for each response type, revealed that for intact pairs and new pairs, there was a main effects of relatedness [intact: *F* (1, 39) = 370.64, *p* < .001, *η*^2^_p_ = 0.91; new: *F* (1, 39) = 25.88, *p* < .001, *η*^2^_p_ = 0.40], and valence [intact: *F* (1, 39) = 22.64, *p* < .001, *η*^2^_p_ = 0.37; new: *F* (1, 39) = 107.67, *p* < .001, *η*^2^_p_ = 0.73], as well as a 2-way interaction between the two factors [intact: *F* (1, 39) = 12.67, *p* = .001, *η*^2^_p_ = 0.25; new: *F* (1, 39) = 7.64, *p* = .009, *η*^2^_p_ = 0.16]. Decomposition of the interaction between relatedness and valence revealed that even though the difference in accuracy rates between neutral and negative pairs emerged for both related and unrelated pairs, it was greater in the latter [intact: *t* _related_(39) = 2.35, *p* = .024, d = 0.37; *t _un_*_related_(39) = 5.26, *p* < .001, d = 0.83; new: *t* _related_(39) = 7.57, *p* < .001, d = 1.20; *t _un_*_related_(39) = 9.97, *p* < .001, d = 1.58]. For rearranged pairs, this analysis revealed main effects of relatedness, *F* (1, 39) = 12.88, *p* = .001, *η*^2^_p_ = 0.25, and valence, *F* (1, 39) = 6.59, *p* = .014, *η*^2^_p_ = 0.15.

(3) **Analyses of item memory ERP effects**

Both rearranged and new pairs do not contain studied associative information. However, while rearranged pairs contain studied items, new pairs contain novel items that were not presented during the study phase. Therefore, the rearranged/new effect, i.e., differences between ERPs associated with correct “rearranged” judgments vs. correct “new” judgments, is indicative of item recognition (e.g., Li et al., 2017; Rhodes & Donaldson, 2008; Zheng et al., 2015). To analyse the item memory effect, we used the same approach that was used for the associative memory effect (in terms of time windows, selected channels, etc., see main text), but included “rearranged” and “new” as the levels of the response type factor. Thus, repeated measures ANOVA was conducted separately for each time window and included four within-subjects factors: relatedness (related or unrelated), valence (negative or neutral), response type (rearranged or new).

**Results**

The topographical distribution of the item memory effect in the various experimental conditions are shown in Supp Fig 1 (for Experiment 1) and 2 (For Experiment 2). The waveforms obtained for new pairs are shown in the main text (Figures 2a and 3a).

Experiment 1

In the early time window (300-550 ms), the ANOVA revealed a 2-way interaction between response type and location, *F* (1.38, 56.65) = 8.49, *p* = .002, *η^2^_p_* = 0.17, and a 3-way interaction between relatedness, valence, and response type, *F* (1, 41) = 6.25, *p* = .017, *η^2^_p_* = 0.13. To decompose the 3-way interaction a follow-up ANOVA with the factors of relatedness and response type was conducted separately for each valence. For negative pairs, this analysis revealed no main effects or interactions that included the response type factor (*ps* > .05). A similar follow-up ANOVA for neutral pairs revealed a main effect of response type, *F* (1, 41) = 6.48, *p* = .015, *η*^2^_p_ = 0.14. Thus, in the early time window, the item memory effect was reliably observed for neutral pairs, but not for negative pairs.

In the late time window (550-800 ms) the ANOVA revealed a main effect of response type, F (1, 41) = 6.77, p = .013, η2p = 0.14, and a 3-way interaction between relatedness, valence, and response type, F (1, 41) = 9.12, p = .004, η2p = 0.18. Decomposition of the 3-way interaction revealed that for negative pairs, there was an interaction between relatedness and response type, F (1, 41) = 6.98, p = .012, η2p = 0.15, depicting a reversed item memory effect (i.e., greater positivity for new vs. rearranged pairs) for related pairs, t (41) = 2.41, p =.021, d = 0.37, but not for unrelated pairs (p = .2). For neutral pairs, a reversed item memory effect was observed, evident as a main effect of response type, F (1, 41) = 7.01, p = .011, η2p = 0.15, with no interaction with relatedness. Thus, overall, in the late time window, a reversed item memory effect emerged in all experimental conditions, except for unrelated negative pairs.

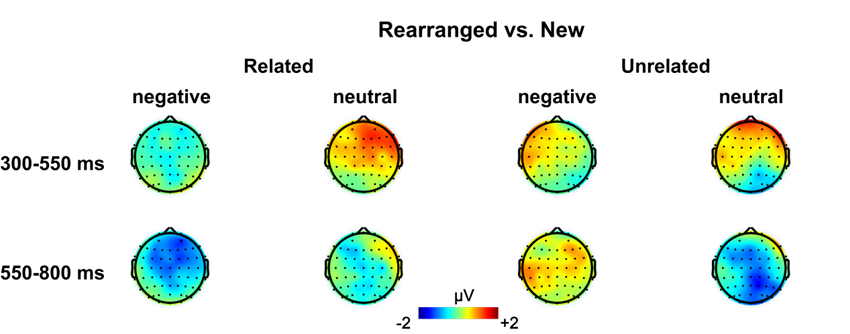

**Supp Fig 1.** Topographical maps of the item memory effects in Experiment 1 in each time window.

Experiment 2

In the early time window (300-550 ms), the analysis revealed a main effect of response type, with more positive-going waveforms for rearranged pairs (vs. new), *F* (1, 39) = 4.55, *p* = .039, *η*^2^_p_ = 0.10, 2-way interaction between relatedness and response type, *F* (1, 39) = 4.17, *p* = .048, *η*^2^_p_ = 0.10, a 2-way interaction between response type and location, *F* (1.25, 48.55) = 10.76, *p* = .001, *η*^2^_p_ = 0.22, and a 3-way interaction between relatedness, valence, and response type, *F* (1, 39) = 9.26, *p* = .004, *η*^2^_p_ = 0.19.

To decompose the 3-way interaction, a follow-up ANOVA with the factors of relatedness and response type was then conducted separately for each valence. For negative pairs, this analysis revealed an interaction between relatedness and response type, *F* (1, 39) = 12.57, *p* = .001, *η*^2^_p_ = 0.24, with a significant item memory effect (rearranged vs. new) for unrelated pairs, *t* (39) = 3.48, *p* =.001, d = 0.55, but not for related pairs (*p* = .36). A similar follow-up ANOVA for neutral pairs revealed no main effects or interactions (*ps* > .05).

In the late time window (550-800 ms), the analysis revealed 2-way interactions between relatedness and response type, *F* (1, 39) = 6.18, *p* = .017, *η^2^_p_* = 0.14, and between valence and response type, *F* (1, 39) = 14.14, *p* = .001, *η^2^_p_* = 0.27. Finally, the analysis revealed a 3-way interaction between relatedness, valence, and response type, *F* (1, 39) = 4.51, *p* = .040, *η*^2^_p_ = 0.10. To decompose the 3-way interaction, we conducted follow-up ANOVAs with relatedness and response type as within-subject factors, separately for each valence. For negative pairs, this analysis revealed a 2-way interaction between relatedness and response type, *F* (1, 39) = 12.81, *p* = .001, *η^2^_p_* = 0.25, resulting from an item memory effect (rearranged vs. new) for unrelated pairs, *t* (39) = 3.45, *p* = .001, d = 0.54 but not for related pairs (*p* = .24). A similar follow-up analysis for neutral pairs revealed a main effect for response type, *F* (1, 39) = 14.19, *p* = .001, *η^2^_p_* = 0.27, with more positive-going waveforms for new vs. rearranged pairs (a “reversed” item memory effect), regardless their relatedness. Thus overall, at the late time window, an item memory effect emerged for unrelated negative pairs, and a reversed item memory effect (of reversed polarity) emerged for related and unrelated neutral pairs.

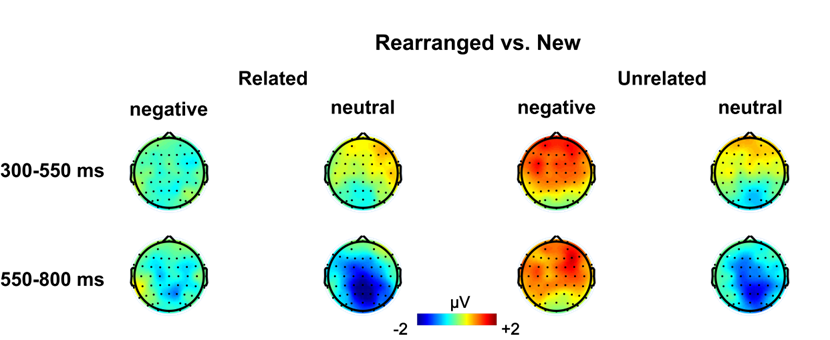

**Supp Fig 2.** Topographical maps of the item memory effects in Experiment 2 in each time window.

**Discussion**

In addition to the associative memory effects reported in the main text, our study showed task-depended modulation of the early rearranged/new effect, which is indicative of item familiarity. Specifically, in Experiment 1, the early rearranged/new effect was greater for neutral (vs. negative) pairs, regardless their semantic relations. In Experiment 2, it was evident for negative unrelated pairs. It is interesting to note that, to some degree, these item memory effects mirror the early associative memory effects, which were obtained for negative pairs in Experiment 1, and for related pairs in Experiment 2. This pattern might indirectly support the suggestion that a shared emotional context and pre-existing semantic relations can promote unitization. It has been suggested that once multiple items have been unitized to create a single representation, only the unitized whole, and not the constituent elements, is readily available for familiarity at retrieval. Consequently, unitization should entail measurable “costs” on item familiarity (Liu et al., 2020; Mayes et al., 2007; Pilgrim et al., 2012). Such costs are evident in our current data, where item-related modulation of the early ERP component was greater for non-unitizable pairs. This suggestion warrants further investigation in future studies.

In the late time window, greater positivity for new pairs (at a magnitude that is equivalent or greater than that of the intact pairs) was observed for related pairs and for unrelated-neutral pairs in Experiment 1, and for neutral pairs in Experiment 2. This somewhat unexpected pattern of intact = new > rearranged seems to be inconsistent with some previous reports of decreased positivity for new vs. intact/rearranged pairs (e.g., Bader et al., 2010; Greve et al., 2007; Kriukova et al., 2013; Mollison & Curran, 2012; Opitz, 2010; Rhodes & Donaldson, 2007, 2008). Nonetheless, it is comparable with other previous findings. For example, Tibon, Gronau, Scheuplein, Mecklinger, & Levy (2014) found a pattern of intact = new > rearranged for unrelated pairs in the late time window; Lu, Liu, Wang, & Guo (2020) found a similar ERP pattern for word-picture pairs in an interactive imagery condition; Addante, Ranganath, & Yonelinas (2012) showed that the late parietal ERP was less positive for low confidence item recognition accompanied by accurate source recognition than for correctly rejected foils. One possibility is that the neural response that we have observed at the late time window reflects a combination of multiple ERP components with overlapping durations. In particular, the late posterior negativity component (LPN) tends to overlap with the traditional late old/new effect, and is of opposite polarity (for a review see Mecklinger et al., 2016). This component is often linked to reconstructive processing, evaluation of retrieval outcomes, and/or task difficulty, although the precise functional role of this old/new difference remains a matter of debate (see, for example, Friedman et al., 2005; Johansson & Mecklinger, 2003; Mecklinger et al., 2016; Sommer et al., 2018; but also recent contradicting results by Park & Donaldson, 2019). In the case of our study, we note that it remains unclear why certain experimental conditions, but not others, would elicit this effect.

(4) **Exploratory analysis: 800-1,000-ms window**

Following a visual inspection of the waveforms, we observed an additional associative memory effect between 800 and 1,000 ms, later than our two pre-defined time-windows, which was apparent in both experiments. As an exploratory analysis, we investigated whether this effect is modulated by relatedness and/or valence.

For Experiment 1, a repeated measures ANOVA with relatedness, valence, response type, and location as within-subject factors, revealed a main effect of response, *F* (1, 41) = 32.36, *p* < .001, *η^2^_p_* = 0.44, a 2-way interaction between response type and location, *F* (1.56, 63.79) = 8.02, *p* = .002, *η^2^_p_* = 0.16, as well as a 3-way interaction between relatedness, response type and location, *F* (1.50, 61.59) = 7.43, *p* = .003, *η^2^_p_* = 0.15. A follow-up ANOVA with the factors of relatedness and response type was then conducted separately for each location. This analysis revealed a main effect of response type in all locations (all *p*s < .001). For parietal locations, this analysis also revealed an interaction between relatedness and response type, *F* (1, 41) = 4.52, *p* = .040, *η^2^_p_* = 0.10, depicting a significant associative memory effect for related pairs, *t* (41) = 3.89, *p* <.001, *d* = 0.60, but not for unrelated pairs, *t* (41) = 1.39, *p* = 0.18, *d* = 0.21.

For Experiment 2, a repeated measures ANOVA at this time window revealed a main effect of response type, *F* (1, 39) = 30.55, *p* < .001, *η^2^_p_* = 0.44, a 2-way interaction between relatedness and response type, *F* (1, 39) = 11.13, *p* = .002, *η^2^_p_* = 0.22, a 2-way interaction between response type and location, *F* (1.61, 62.77) = 7.86, *p* = .002, *η^2^_p_* = 0.17, and a 3-way interaction between relatedness, response type, and location, *F* (1.64, 63.78) = 3.69, *p* = .039, *η^2^_p_* = 0.09. A follow-up ANOVA with the factors of relatedness and response type was then conducted separately for each location. This analysis revealed main effects of response type in all locations (all *p*s < .001), as well as a 2-way interaction between relatedness and response type in central and parietal locations [central: *F* (1, 39) = 9.95, *p* = .003, *η^2^_p_* = 0.20; parietal: *F* (1, 39) = 13.57, *p* = .001, *η^2^_p_* = 0.26]. Decomposition of this interaction revealed that in central locations, the associative memory effect was greater for related pairs, *t* (39) = 5.23, *p* < .001, d = 0.83, vs. unrelated pairs *t* (39) = 2.61, *p* = .013, d = 0.41. In parietal locations the effect was only observed for related pairs, *t* (39) = 5.44, *p* < .001, d = 0.86, but no for unrelated pair, *t* (39) = 1.34, *p* = .19, d = 0.21.

These exploratory analyses revealed an additional associative memory effect, which was widespread at a later time window ranging from 800-1,000 ms post stimulus onset. At parietal locations, this effect was further modulated by semantic relatedness, and thus corresponds to the behavioral advantage observed for related (vs. unrelated) pairs. Parietal regions had been implemented in sustained recollective processes that track the period over which recollected information is maintained (e.g., Vilberg & Rugg, 2008; Vilberg & Rugg, 2012; Vilberg & Rugg, 2013; see also Humphreys et al., 2021, for a unifying cross-domain account). The elaborated encoding afforded by preexisting semantic relations might be gradually elicited, and therefore reflected in a later time window than our predefined one. We thus suspect that this late modulation reflects the continuous involvement of recollective processes (that is, an extension of the activation observed in the previous time window).
